# Hornbill abundance and habitat correlates in Kali Tiger Reserve, Western Ghats, India—Insights from collaborative monitoring

**DOI:** 10.64898/2026.04.18.719330

**Authors:** Nandita Madhu, Himanshu Lad, Bindu Kempegowda, Vishal Sadekar, Sharvani Deshpande, Nitin Kawthankar, Navendu Page, Ashwin Bhat, Nilesh Shinde, Rohit Naniwadekar

## Abstract

The future of species, particularly large-bodied birds, under climate threats, is increasingly unclear. Protected areas (PAs) can mitigate the impacts of global change, but are not completely immune. This makes it imperative to monitor populations even inside PAs. In the forests of the Western Ghats, three hornbill species are found—Malabar Grey Hornbill, Malabar Pied Hornbill and Great Hornbill. Kali Tiger Reserve, of Karnataka state, India, is known for its healthy hornbill populations, yet few systematic surveys have attempted to estimate their abundance, distributions and habitat affiliations. In this study, we documented the distributions of the three hornbills, estimated the density of Malabar Grey Hornbills and assessed correlates of hornbill encounter rates within the reserve. Malabar Grey Hornbills were the most abundant and widely distributed, found in all ranges of the reserve. Gund, Kadra and Phansoli had the most detections of Malabar Pied and Great Hornbills. Encounter rates of Malabar Grey Hornbill were positively correlated with food-tree stem density, those of Malabar Pied Hornbill negatively correlated with basal area, while those of Great Hornbill not significantly associated with any variable. Malabar Grey Hornbills had a density of 5.1 per km^2^ (mean flock size = 1.2). While encounter rates of Malabar Pied and Great Hornbills were low, these numbers track the breeding season, when vocal activity is low and females are inside nests. This survey represents a partnership between researchers and the Karnataka forest department, aimed towards inculcating the collaborative spirit of research and monitoring.

## 2. INTRODUCTION

Tropical forests host more than two-thirds of the world’s biodiversity, also storing a quarter of the world’s terrestrial above and belowground carbon. However, they only cover about 10% of the Earth’s land surface (Bonan, 2008; Lewis, 2005) and are globally threatened by deforestation, degradation, over-harvesting, invasion and climate change, anthropogenic or otherwise (Gardner et al., 2009). Not only are they hubs of diversity but also of extinctions. The future of wildlife has never been more uncertain (Bradshaw et al., 2009). Due to this uncertainty, it becomes essential to estimate and monitor wildlife populations across time and space. Large-bodied forest specialist species are a suitable candidate to monitor trends in populations since they require large relatively intact forest patches, occur in low densities and have slow breeding rates.

Today, Protected Areas (PAs) are the most widely used system for preserving flora, fauna and ecosystem services worldwide and will probably remain so (Joppa & Pfaff, 2011). They have been demonstrated to reduce tree felling (Andam et al., 2008), urbanisation and agricultural expansion (Moraes et al., 2017), also providing buffers against climate change (Xu et al., 2022). They are not completely immune to climate warming and species could be negatively impacted by rising temperatures but losses in some PAs can be offset by increases in others (Thomas & Gillingham, 2015). However, all areas cannot be predicted to have the same outcomes which is why it is necessary to estimate and monitor wildlife populations even inside PAs.

Hornbills are large-bodied species known for their ecological role in seed dispersal, facilitating forest regeneration. They have large home ranges and Asian species can travel long distances, up to 13 km, to deposit seeds (Naniwadekar et al., 2019). They are secondary cavity nesters, making use of pre-existent tree cavities in large trees (Poonswad, 1995). Hence, it is crucial to have large, undegraded trees with hollows to host healthy hornbill populations. They are also dependent on fruiting resources and are known to track fruit availability or food-tree density across space and time (Kinnaird & O’Brien 1996, Naniwadekar et al., 2015, Pawar et al., 2021).

The Western Ghats, a mountain chain in the southern part of India, with large expanses of tropical forest vegetation and spanning over 1400 km, is one of the country’s four global biodiversity hotspots (Myers et al., 2000). Spread across six states, its evergreen and moist deciduous forests provide vital habitats for avian diversity. Notably, three species of hornbills - Malabar Grey Hornbill (*Ocyceros griseus*), Malabar Pied Hornbill (*Anthracoceros coronatus*) and Great Hornbill (*Buceros bicornis*) inhabit the region. Malabar Grey Hornbills inhabit the wet forests of the Western Ghats, from Nashik, Maharashtra to the Kanniyakumari and Agasthyamalai hills in the south. The Malabar Pied and Great Hornbills are more widespread, with the former occupying the deciduous and open areas of central and eastern India, the Western Ghats and Sri Lanka and the latter occupying large tracts of primary evergreen forests of the Western Ghats, Himalayan foothills, and countries in Southeast Asia.

Hornbills are threatened by habitat loss, degradation, fragmentation, hunting pressures, and loss of nesting trees (Naniwadekar et al., 2015; Shankar Raman & Mudappa, 2003; Sriprasertsil et al., 2024). Given the requirements of some species such as the Great Hornbill (*Buceros bicornis*) for large tracts of less disturbed forests, PAs are often refuges for their survival. Moreover, climate and weather impact their distributions and nesting success (van de Ven et al., 2020, Sarkar & Talukdar, 2023). Given that hornbills are slow-breeders, reaching sexual maturity after 2-4 years with a clutch size of 1-3 a year, it is vital to monitor their populations. A recent study found declines in Malabar Grey Hornbill (*Ocyceros griseus*) populations from within a PA in the absence of contemporary perceivable threats (Pawar et al., 2021). This necessitates periodic monitoring of hornbills inside PAs, which will help in detecting changes in populations and implementing timely interventions for their conservation. One of the major gaps is the absence of baseline information on their abundances even from inside PAs.

Kali Tiger Reserve (KTR) is a PA situated in Uttara Kannada, the district with the highest green cover in the Western Ghats (Ramachandra et al., 2016). It is thought to host substantial populations of hornbill species, especially those of Malabar Grey and Malabar Pied Hornbills (Mudappa & Shankar Raman, 2009). Hence, there is a need for monitoring abundances, habitat associations and threats across the reserve for effective management of hornbill populations. While previous studies have demonstrated the utility of line transects for accurately estimating hornbill densities, it is imperative to have the transects spread out uniformly across the focal study region to accurately estimate their populations. As part of the annual monitoring of tiger prey, staff across Tiger Reserves mark and monitor line transects that are almost uniformly spread out across the entire Tiger Reserve. These transects are 1-2 km long and ideal for estimating and monitoring hornbill populations. Assisted by forest staff, across seven ranges of the reserve for the duration of the breeding season (March-May), we aimed to determine the 1) distributions of Malabar Grey Hornbill, Malabar Pied Hornbill and Great Hornbill in KTR, 2) correlates of hornbill encounter rates in KTR, and 3) densities of Malabar Grey Hornbill in the ranges of KTR.

## 3. MATERIALS AND METHODS

### 3.1. Study Area

KTR, spread over an area of 1,425 km^2^, lies between 14° 57’ 23.04” N; 74° 15’ 7.56” E and 15°9’56.16” N; 74°43’ 10.56” E in Uttara Kannada district of Karnataka. The elevation in KTR ranges from 27–1059 m ASL. Rainfall received varies from 1,250 mm towards the eastern side to about 5,000 mm towards the western part of KTR, which forms the crestline of the Sahyadri. KTR harbours a diverse array of habitats, including, South Indian Moist Deciduous Teak Forests (3B/C1), Southern Moist Mixed Deciduous Forests (3B/C2), West Coast Semi-Evergreen Forests (2A/C2), Moist Bamboo Brakes (2B/E3) and Cane Brakes (2B/E1) (Champion and Seth, 1968; Tripathy et al., 2024). KTR is one of five tiger reserves of Karnataka and has an area of 1,425 km^2^. It harbours tigers (*Panthera tigris*), leopards (*Panthera pardus*), wild dogs (*Cuon alpinus*), gaur (*Bos gaurus*), sloth bears, a small population of elephants (*Elephas maximus*), among other wildlife. Along with Bhimgad Wildlife Sanctuary to the north, and adjoining Protected Areas in Goa, it is a critical stronghold for hornbills in Western Ghats. This study was conducted in selected locations across the reserve, covering a range of habitats from evergreen and deciduous forests (Fig. 1). Evergreen and semi-evergreen forests house large trees such as *Tetrameles nudiflora* and *Terminalia bellirica*, that are suitable for hornbill cavities. Moreover, they also harbour a diverse array of flesh-fruited plants such as *Ficus amplissima*, and culturally and ecologically significant species such as *Myristica* spp., *Knema attenuata*, and *Canarium strictum* which are important hornbill food plants. A human population of more than 29,000 (∼6,900 families) mostly belonging to Kunbi, Maratha, and Gouli communities reside within KTR.

**Figure 1.**
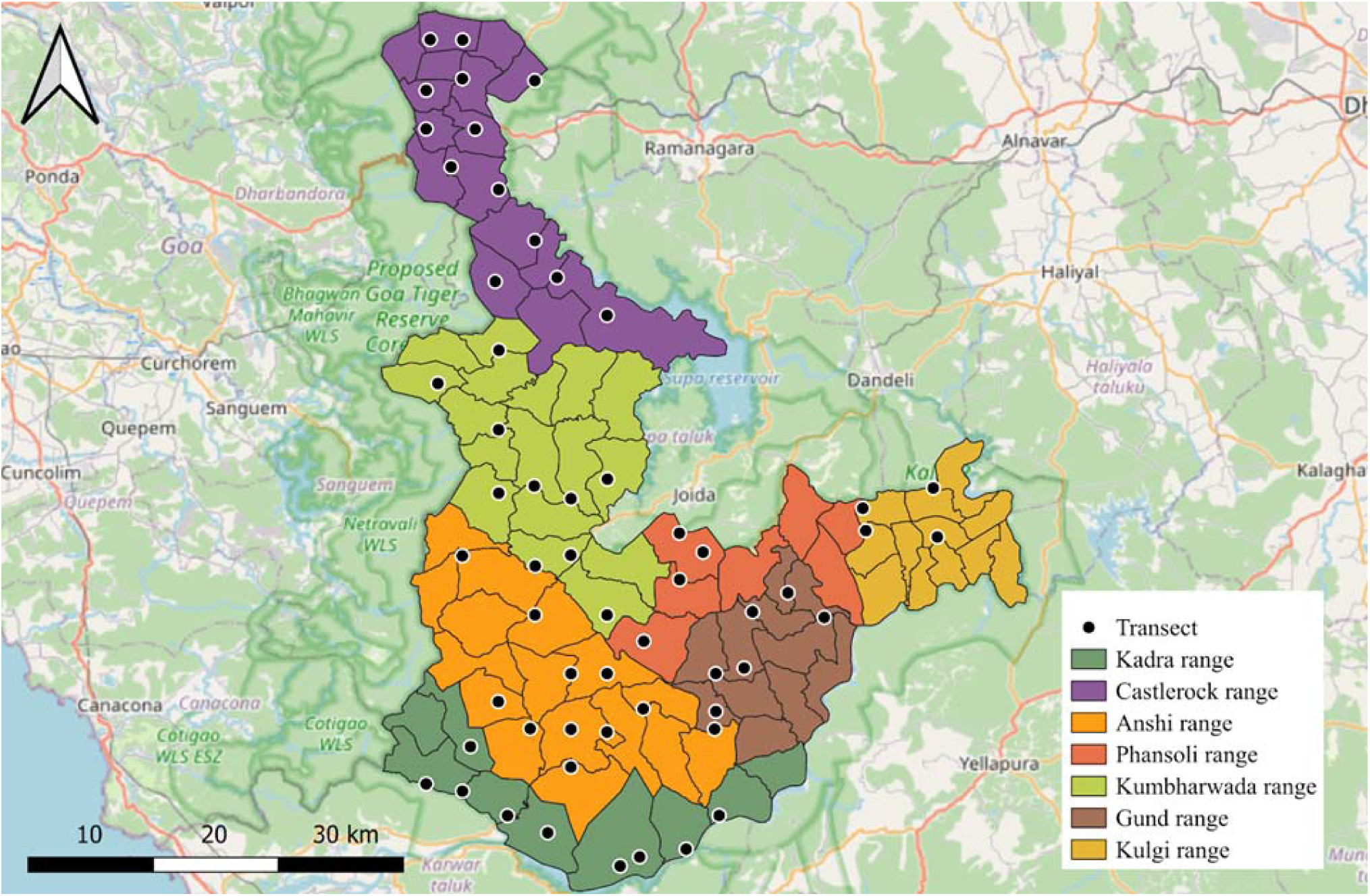
Map, displaying the 57 transect locations sampled across the seven ranges of the reserve.

### 3.2. Field Methods

#### 3.2.1. Bird sampling

We sampled 57 transects laid for monitoring tiger prey across the reserve (Fig.1, Table S1). The starting points of the transects are randomly chosen, while ensuring that each beat has at least one transect. The transects are laid in N-S orientation. We walked each transect once between March and May 2025, coinciding with the breeding season of hornbills. Two observers (NM and HL) assisted by forest department staff, sampled birds by walking in a slow, steady pace, noting down species identity, whether the individual was seen or heard, flock size, age-sex ratio (when detected visually), perpendicular distance (when detected visually), and distance-band (when detected auditorily). Distance-bands were set in the following intervals (m): 0–2, 3–5, 6–8, 9–11, 12–15, 16–20, 21–25, 26–30, 31–50, 51–70, 71–100, 101–150, 151–200 and > 200. We observed birds in the morning between 0600–1000 hrs. We also recorded hornbill occurrences opportunistically outside transect surveys.

#### 3.2.2. Vegetation sampling

We laid fifty six (50×10 m^2^) plots (Fig. 2) distributed across the deciduous and evergreen forests of the reserve. In each plot, we recorded the species identity, girth and height of all woody stems ≥ 10 cm girth at breast height (GBH). We estimated stem densities of all species, including those that are food plants for hornbills (Table S2, Ahirbudhnyan et al., 2026), basal area and canopy height.

**Figure 2.**
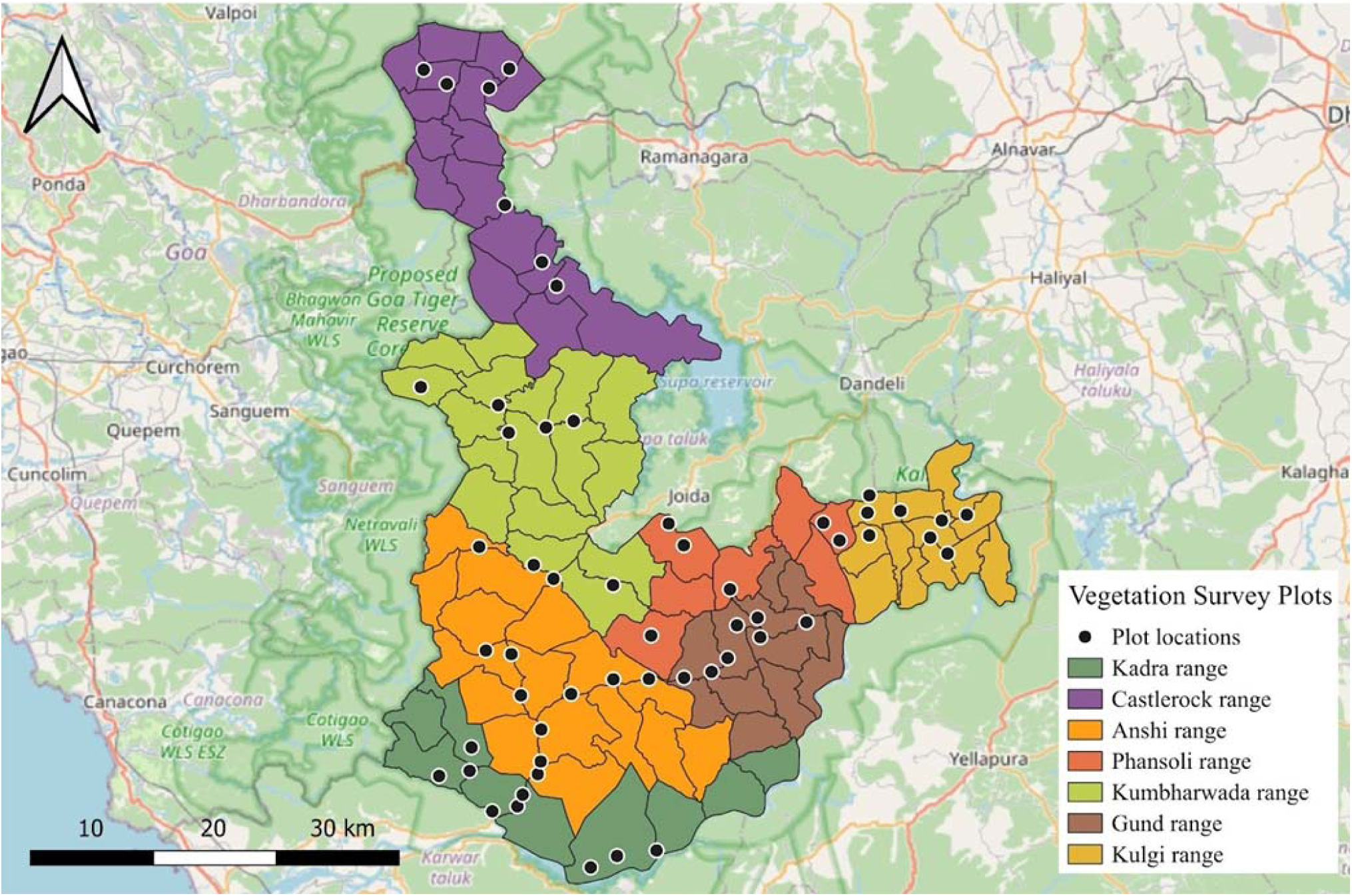
Map, displaying the 57 vegetation plot locations sampled across the seven ranges of the reserve.

### 3.3. Analysis

We conducted all the analyses in R version 4.4.2 (R Core Team, 2024).

#### 3.3.1. Distributions of Malabar Grey Hornbill, Malabar Pied Hornbill and Great Hornbill in KTR

Distributions were recorded using the application “Epicollect5” (v. 87.2.3) by recording the geographical coordinates of the location of every detection, opportunistic or otherwise. Distribution maps were generated using QGIS.

#### 3.3.2. Correlates of Hornbill encounter rates in the ranges of KTR

We estimated encounter rates (per km) of the three hornbill species. We checked for correlations between the encounter rates of Malabar Grey Hornbill, Malabar Pied Hornbill and Great Hornbill and variables—stem density (ha^-1^), food-tree stem density (ha^-1^), canopy height (m) and basal area (m^2^ ha^-1^). These vegetation covariates are associated with forest structure (stem density, canopy height, and basal area) and food availability. Hornbills have been shown to demonstrate affiliations to structural variables that can indicate vegetation type, and resource variables such as fruit-tree density (Dasgupta et al., 2022; Shankar Raman & Mudappa, 2003). We used Pearson’s correlation coefficients (since variables were normally distributed) and coefficients for which *p* < 0.05 were considered significant.

#### 3.3.3. Densities of Malabar Grey Hornbill in the ranges of KTR

We estimated the abundance and density of Malabar Grey Hornbill using the multiple covariate distance sampling (MCDS) approach in the R-package ‘Distance’ (version 2.0.0, Miller et al. 2019). We modelled the detection probability as a function of forest ranges, since the ranges coincide with the broad distribution of habitats (Evergreen: Castlerock, Kumbharwada, Anshi; Deciduous: Kadra, Phansoli, Gund, Kulgi) (Kempegowda et al., in prep.), which can influence detection probability. We truncated detections beyond 150 m from the point of observation to improve model fit. We fitted half-normal and hazard-rate detection function models and selected the best-fitting model based on the lowest Akaike Information Criterion (AIC) value (Buckland et al., 2015) and Cramer-von-Mises goodness of fit test. Apart from overall densities, we also estimated densities separately for every range of KTR.

## 4. RESULTS

### 4.1. Distributions of Malabar Grey Hornbill, Malabar Pied Hornbill and Great Hornbill in KTR

A majority of our sightings were of Malabar Grey Hornbill (Table S3). We detected them in all ranges of the reserve. Kumbarwada was a hotspot of Malabar Grey Hornbill sightings.

Malabar Pied Hornbills were relatively rare across the landscape, with sparse sightings in Castlerock and Anshi, and almost none in Kumbarwada (Fig. 3b). Sightings peaked in Kadra and Gund. We saw large flocks near river Kali in Kadra, one of which had multiple females but only two males. They were also seen utilising mixed forests and plantations. We observed one Malabar Pied Hornbill foraging on the fruits of *Acacia auriculiformis* (a non-native species that is often planted).

**Figure 3.**
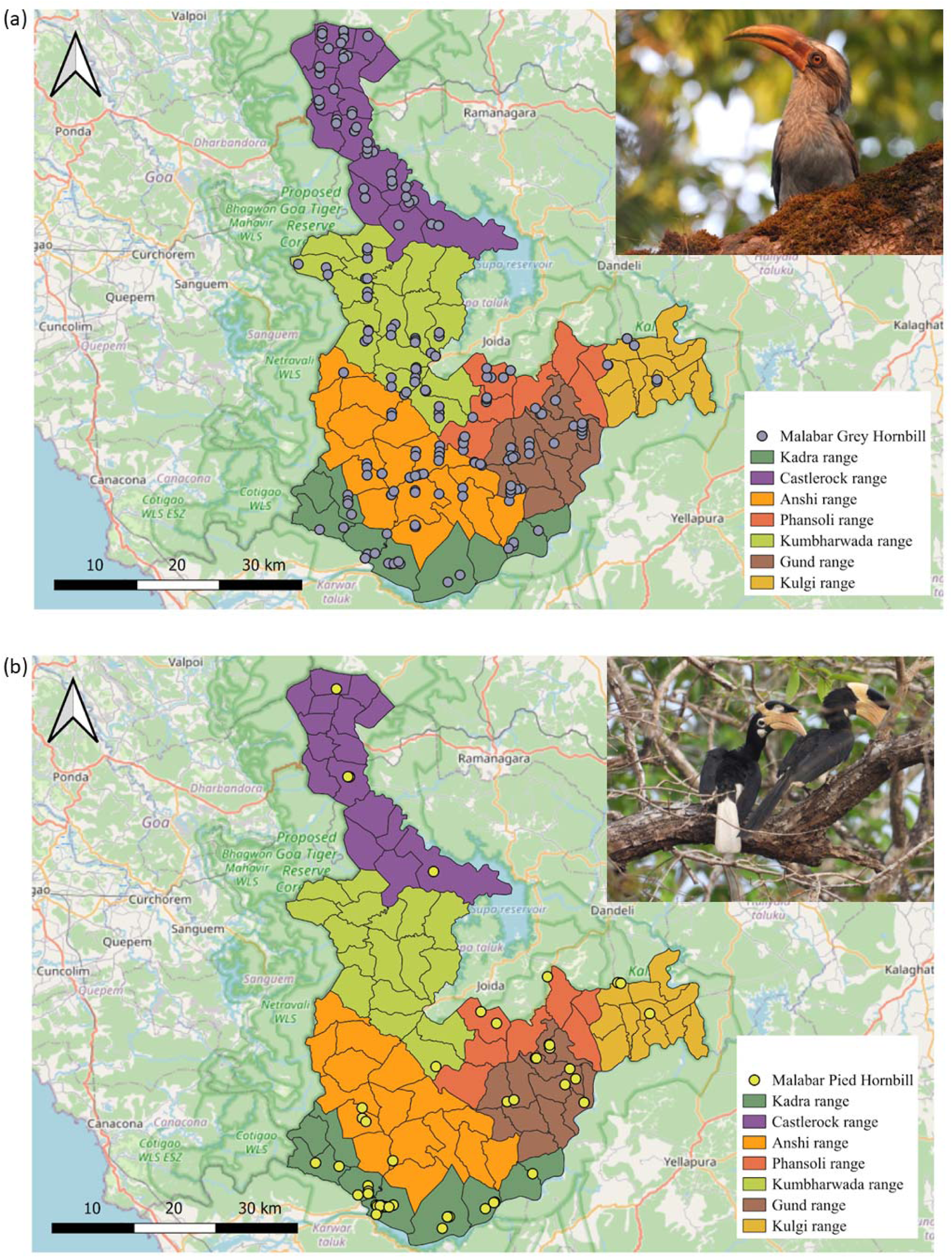
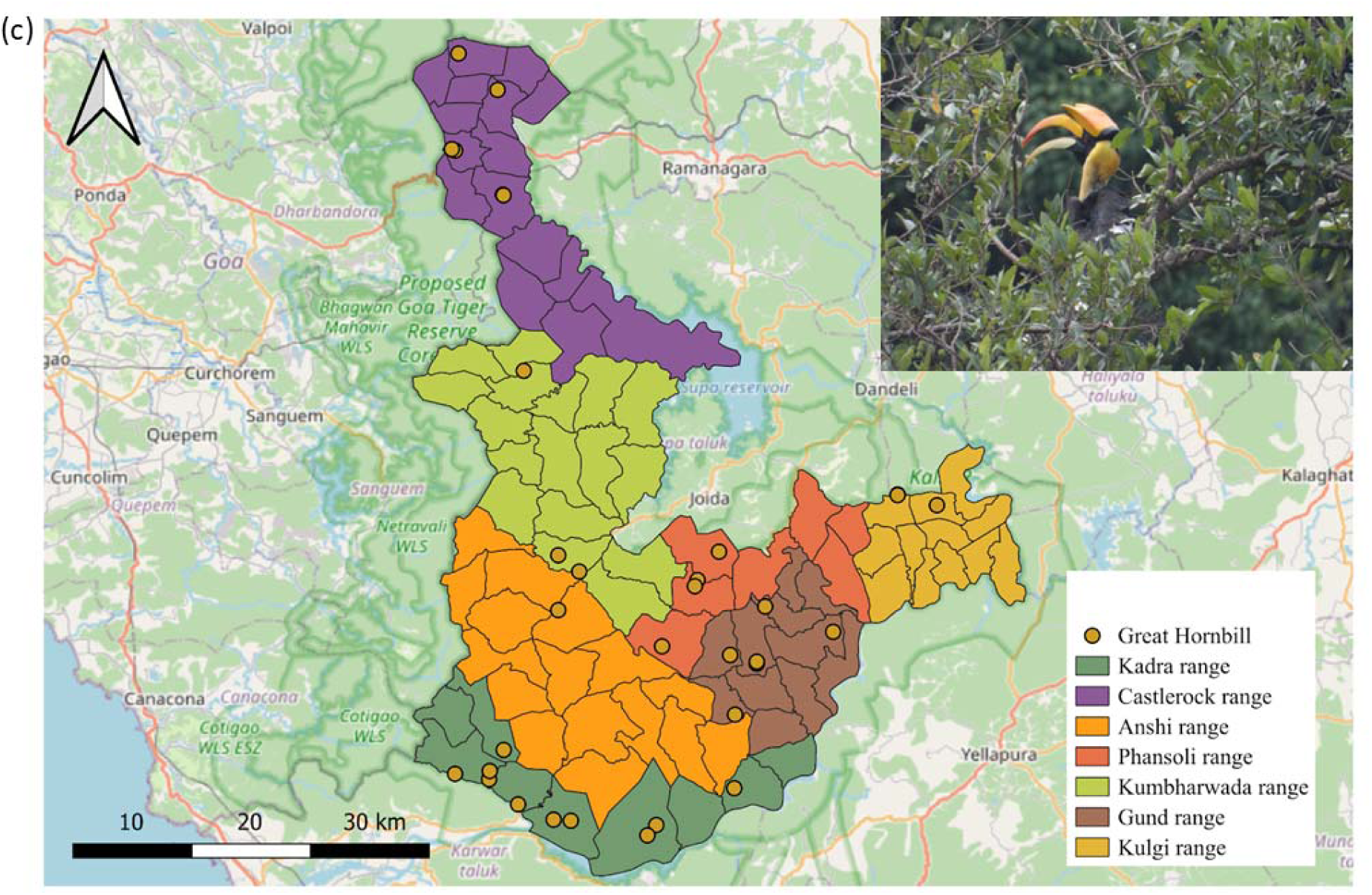
Detections of (a) Malabar Grey Hornbill *Ocyceros griseus*, (b) Malabar Pied Hornbill *Anthracoceros coronatus* and (c) Great Hornbill *Buceros bicornis* across Kali tiger reserve. Detections include visual and aural detections during and outside transects when detected opportunistically.

Great Hornbills, like the Malabar Pied Hornbills, were also rare within the reserve. We only had one detection in Anshi and two in Kulgi. Most detections were of calls.

### 4.2. Correlates of Hornbill encounter rates in the ranges of KTR

Overall encounter rates across the reserve of Malabar Grey Hornbill, Malabar Pied Hornbill and Great Hornbill were 1.4 (± 0.1 SE) km^-1^, 0.1 (± 0 SE) km^-1^, and 0.2 (± 0.1 SE) km^-1^, respectively (Table 1). Encounter rates of Malabar Grey Hornbill were positively correlated with food-tree stem density (Fig. 4a, Table S5a). Encounter rates of Malabar Pied Hornbill were negatively correlated with basal area (Fig. 4b, Table S5b), while those of Great Hornbill were not significantly associated with any variable (Table S5c).

**Figure 4.**
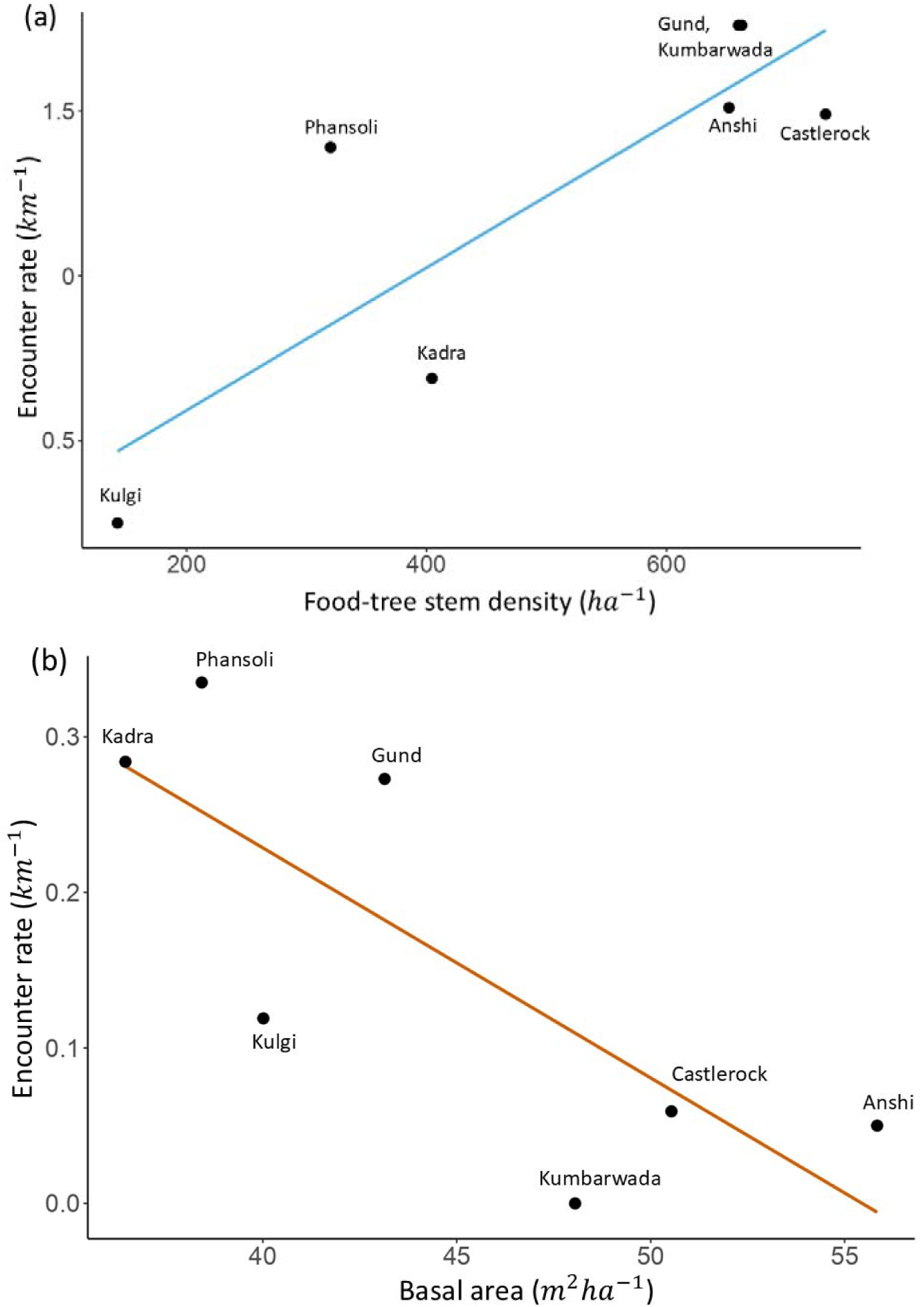
Scatterplots showing the significant associations (*p* < 0.05, for Pearson’s correlations) of the encounter rates of (a) Malabar Grey Hornbill *Ocyceros griseus* with food-tree stem density (ha^-1^) and (b) Malabar Pied Hornbill *Anthracoceros coronatus* with basal area (m^2^ ha^-1^).

**Table 1.**
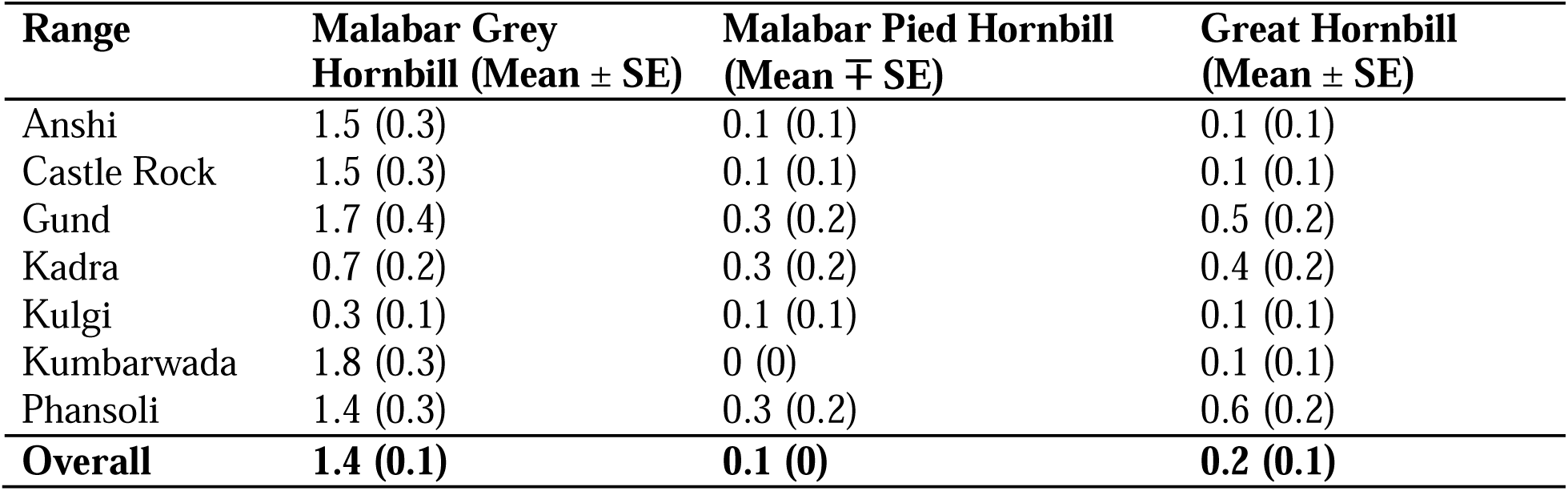
Overall and range-wise flock encounter rates (km^-1^) of Malabar Grey Hornbill, Malabar Pied Hornbill and Great Hornbill in KTR.

### 4.3. Densities of Malabar Grey Hornbill in the ranges of KTR

We found that the half-normal model with no covariate (Table S4) had the least AIC value, indicating that detection probability did not differ significantly between ranges (Table S4). Overall flock size of Malabar Grey Hornbills as estimated using only the visual detections was 1.2 (± 0.1 SE) birds, and the overall detection probability (*p*) was 0.8 (± 0.1 SE). Density estimates for the species ranged between 1.1–7.8. km^-2^, in which the highest value was in Gund and the lowest in Kulgi. Range-wise abundance estimates ranged from 123–2191 birds. Density and abundance estimates are presented in Table 2.

**Table 2.**
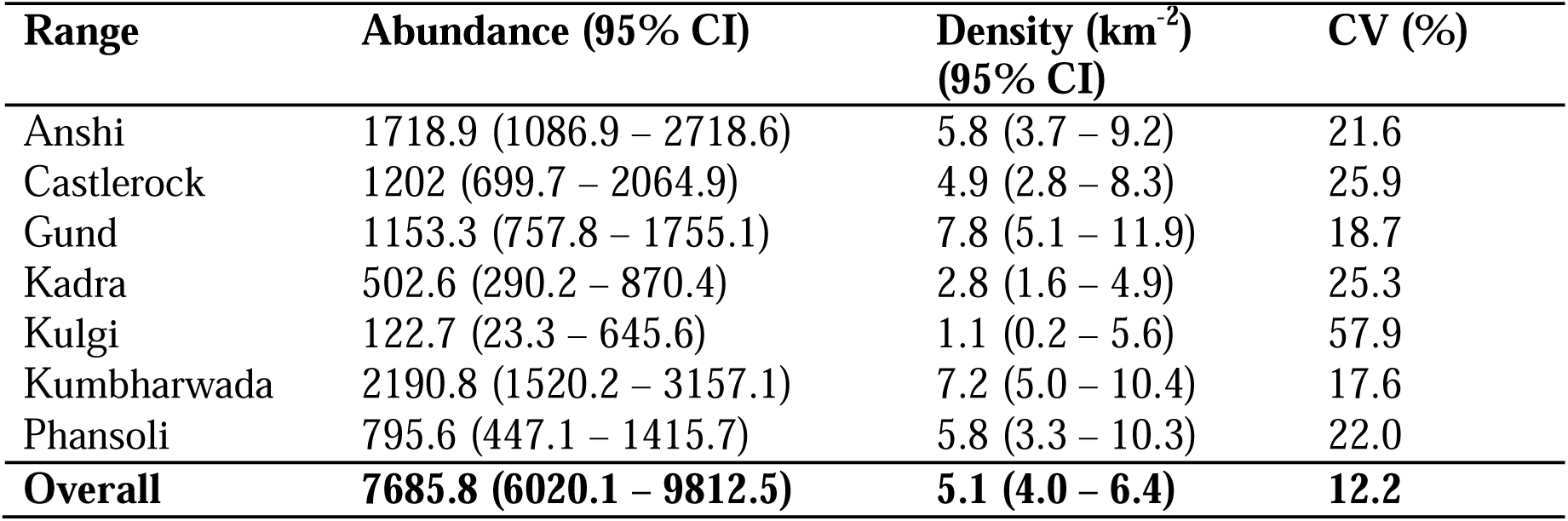
Overall and range-wise densities of Malabar Grey Hornbill in KTR.

## 5. DISCUSSION

This is the first systematic survey of hornbills in Kali Tiger Reserve that provides information on distribution and abundance of three of the four hornbills found in the region. The study demonstrates the association of the Malabar Grey Hornbills with wetter habitats and areas with higher food resource abundance, and of the Malabar Pied Hornbills with deciduous habitats in the reserve. The study suggests that there could be at least 6,000 Malabar Grey Hornbills in Kali Tiger Reserve, highlighting the importance of the area for an endemic hornbill species that has experienced population declines in other parts of the region. Such surveys with systematic spatial coverage and replication across large areas are rather limited in the region (David et al., 2019) and across Asia (Pradhan et al., 2024), highlighting the need for such surveys in other important hornbill sites. We also demonstrate the utility of transects established for estimating tiger prey populations for estimating hornbill densities with precision.

### 5.1. Distributions and relative abundance of Malabar Grey Hornbill, Malabar Pied Hornbill and Great Hornbill

While, Malabar Grey Hornbills were present in all ranges of the reserve, consistent with past knowledge of habitat use (Balasubramanian et al., 2007), they were more abundant in evergreen forest patches than in deciduous, with Kulgi Range, that primarily harbours deciduous forests, having the least encounter rate. They were observed in high numbers in Kumbarwada and Gund, both ranges primarily hosting evergreen and semi-evergreen forests (Kempegowda et al., in prep.). The wet forests, which generally have higher stem density and basal area, may also harbour greater diversity and abundance of hornbill food plants. The encounter rates of Malabar Grey Hornbill was positively associated with hornbill food plant density. This is in line with the findings of Shankar Raman & Mudappa, (2003) and Pawar et al., (2021). This also suggests patch-level fruit tracking behaviour of several hornbill species around Asia (Kinnaird & O’Brien 1996, Anggraini et al., 2000).

Generally, the smaller hornbill species are more abundant than the larger ones (but see Sriprasertsil et al., 2024). Encounter rates of Malabar Grey Hornbills were higher than Great Hornbills, similar to patterns observed by Mudappa & Shankar Raman (2009) and Kannan & James (1999) Malabar Pied Hornbills had the least encounter rate, about ten times lower than Malabar Grey Hornbills. Pawar & Sadekar (2023) in the northern Western Ghats, also found that Malabar Grey Hornbills were more abundant than Malabar Pied Hornbills, though the difference was not as stark. In KTR, they had high encounter rates in Gund, Phansoli and Kadra. Kadra has a reservoir and alongside the river, we detected multiple large flocks. Gund and Phansoli had deciduous forests where we detected the hornbills on transects.

Mudappa & Shankar Raman (2009) found encounter rates ranging from 0-0.6/km across five transects in the Anshi-Dandeli region. Despite KTR and Dandeli Territorial Range situated in landscapes contiguous with each other, they seem to have considerable differences in the encounter rates of this species. Vijayakumar & Davidar (2011) found high encounter rates of this species (4 km^-1^) which was a reduction from an even higher figure (of 8 km^-1^) in 1983–1984 (Reddy et al., 1990). They sampled five areas, among which only one overlapped with ours (Kulgi) and laid transects in reserved forests that fall outside the PA network. Their spatial coverage was low compared to the one in this study (7.5 km, spanning six transects) and restricted to the moist deciduous forests, which are the preferred habitats, particularly of Malabar Pied Hornbill. Therefore, encounter rates may be high in those transects. The Dandeli Territorial Range is generally known for being a stronghold for Malabar Pied Hornbill populations (Datta & Sakthivel, 2010). These differences may have come about as a significant area of KTR has large tracts of evergreen and semi-evergreen forests and most ranges of the reserve have some evergreen cover. Malabar Pied Hornbills are affiliated to moist deciduous forests or riverine areas (Balasubramanian et al., 2007), known to exhibit more activity and also roost in such habitats. On the other hand, existing forest patches in Dandeli Territorial Range are moist deciduous or mixed deciduous, dominated by species such as *Terminalia paniculata*, *Lagerstroemia microcarpa* and *Tectona grandis* (Vijayakumar & Davidar, 2011). Their preference for more deciduous habitats also likely resulted in the negative correlation between Malabar Pied Hornbill encounter rates and basal area, which is typically lower in the deciduous forests compared to the evergreen, which may have higher density and generally larger trees than deciduous forests.

The encounter rate of Great Hornbills in KTR was similar to its overall encounter rate (0.28/km) in wildlife sanctuaries across the Western Ghats (Mudappa & Shankar Raman, 2009). Surprisingly, we did not encounter them much in the evergreen forest stretches as expected. They were documented at higher rates in the transects of Kadra and Phansoli, both of which hosted largely moist deciduous forests. Great Hornbills are known to occur in moist deciduous forests in northern Western Ghats and in the Western Himalaya. In KTR, there could also be variation in detection probability for Great Hornbills across habitat types, though we did not detect differences in detection probability across habitats for Malabar Grey Hornbills. Additional sampling effort in future studies could allow systematic comparisons of hornbill densities across habitat types with different forest structural characteristics. We did not find statistically significant correlations between Great Hornbill encounter rates with forest structure and food plant density. Hornbills may track resources at different spatial scales, and Great Hornbills were found to track trees with large fruit crops instead of patches with higher food plant densities (Kannan & James, 1999, 2008; Naniwadekar et al., 2015). A previous study in southern Western Ghats also failed to find positive correlation between Great Hornbills and ripe fruiting *Ficus* trees (Pawar et al., 2021) suggesting that Great Hornbills may be tracking fruit resources at different spatial scales than that examined in this study.

### 5.2. Densities of Malabar Grey Hornbill in the ranges of KTR

The overall density of Malabar Grey Hornbill (5 individuals per km^2^) across the reserve was low compared to an older estimate from the Amboli-Goa-Dandeli region (9.4 individuals km^-2^) (Mudappa & Shankar Raman, 2009), but similar to that estimated by Pawar & Sadekar., (2023) (5.1 individuals km^-2^) in Tillari region, northern Western Ghats. The spatial coverage of Mudappa & Shankar Raman, (2009) was low compared to this study and might have been restricted to the wetter sites, which might be the reason behind lower estimates in this study. Moreover, we found significant differences in densities of Malabar Grey Hornbill between the dry and wet habitats of KTR. Our estimates of Malabar Grey Hornbills in the wet forests were comparable to those estimated by Mudappa & Shankar Raman, (2009). However, the Malabar Grey Hornbill densities are significantly lower in this region compared to the evergreen forests of Anamalai hills, where the nesting densities of Malabar Grey Hornbills (Mudappa & Raman, 2009: 23.9-33.1 birds km^-2^; Pawar et al., 2021: 17.5 birds km^-2^) were almost twice that of KTR but similar to those estimated from Periyar Tiger Reserve. This is likely because of greater resource availability in Anamalais compared to KTR, an aspect that needs further investigation.

### 5.3. Prospects for future research

This study was conducted in the timespan of a few months (March-May) that coincide largely with the nesting season of Malabar Grey (February-May) and Pied Hornbills (March-May), and partially with that of the Great Hornbill (December-May) (Pawar et al., 2021). This study provides important baselines of hornbill distribution and abundance during the nesting season. Future studies could extensively assess these parameters during nesting and non-nesting seasons to provide comprehensive estimates of densities. The Reserved Forests under the territorial region adjoining KTR should also be systematically surveyed as it has a notable Malabar Pied Hornbill population. These could be supplemented with habitat and resource affinities to variables such as canopy height, basal area, canopy cover, tree density, host-tree density, food-tree density and fruit availability.

This project was undertaken in collaboration with the forest department of the state of Karnataka. The Forest Department staff were closely involved in helping plan, conduct the field surveys, and facilitating logistical support for the researchers. The data and results were shared back with the department, reflecting a partnership between researchers and forest managers. Co-production of knowledge using transects established for tiger prey monitoring greatly facilitated reliable estimation of hornbill abundances at reduced costs. There are 58 other tiger reserves in India, many of which are important hornbill habitats. Such efforts can be potentially replicated at other sites (see David et al., 2019), facilitating reliable estimation of hornbill densities across a diverse range of habitats in the country. Transdisciplinary partnerships with managers can help overcome accessibility and logistical constraints while enabling researchers to draw on the knowledge from managers and field staff, who are mostly indigenous community members (Laborde et al., 2018; Paasche, 2016). Such steps and initiatives, that move toward cultivating a more participatory and collaborative nature of research, are highly recommended and timely in the face of ongoing anthropic and climate-induced impacts on wildlife.

## 7. CRediT AUTHORSHIP CONTRIBUTION STATEMENT

Conceptualisation: RN, NM, HL, BK; Data curation: NM, HL, BK; Formal analysis: NM, RN; Funding acquisition: RN; Investigation: NM, HL, BK, VS, SD, NK, NP, RN; Methodology: RN, HL, NM; Project administration: NS, HL, NM, RN; Resources: NS, RN, AB; Supervision: RN, NS; Visualisation: NM, RN; Writing – original draft: NM; Writing – review & editing: RN, NS, HL, BK, VS, SD, NK, NP, AB. All authors in this study are indigenous to the country where the study was conducted. Whenever relevant, literature published by scientists from the region has been cited.

## 8. ACKNOWLEDGEMENTS

We would like to thank the Karnataka Forest Department for granting us the necessary permission (KFD/WL/E2(RE)/155/2024/1619433) and support to conduct the study—Principal Chief Wildlife Warden (PCCF) and Chief Wildlife Warden (CWLW), Chief Conservator of Forests (Kanara Circle)—Shri. Vasanth Reddy, Assistant Conservator of Forests (ACFs), Range Forest Officers (RFOs) and Forest Department field and office staff. We would also like to thank members of the Wildlife Research and Training Centre (WRTC), Kali Tiger Reserve. We thank Rintu Mandal for help with analyses. We are grateful to Rohini Nilekani Philanthropies and On the Edge for Conservation for providing financial support for this work.

## 10. DATA ACCESSIBILITY

The data and code that support the findings of this study will be made openly available on Zenodo upon acceptance of the manuscript.

## FIGURES AND TABLES

This section contains the following figures and tables cited in the main manuscript:

## SUPPLEMENTARY INFORMATION

This section contains the following information:

### Supplementary tables

**Table S1.** Summary of transects walked, ranges covered, and effort undertaken across Kali Tiger Reserve for hornbill population estimation.

| Transect ID | Range | Length (km) |
| --- | --- | --- |
| T01 | Kulgi | 2 |
| T02 | Kulgi | 2 |
| T03 | Kulgi | 1.8 |
| T04 | Kulgi | 2.1 |
| T05 | Gund | 1.7 |
| T06 | Gund | 1.35 |
| T07 | Gund | 1.8 |
| T08 | Gund | 1.9 |
| T09 | Gund | 2 |
| T10 | Gund | 1.4 |
| T11 | Anshi | 1.8 |
| T12 | Anshi | 1.9 |
| T13 | Anshi | 2 |
| T14 | Anshi | 2 |
| T15 | Anshi | 1.9 |
| T16 | Anshi | 2 |
| T17 | Anshi | 2.1 |
| T18 | Anshi | 2 |
| T19 | Anshi | 2 |
| T20 | Anshi | 1.9 |
| T21 | Kadra | 1.9 |
| T22 | Kadra | 2 |
| T23 | Kadra | 1.9 |
| T24 | Kadra | 1.8 |
| T25 | Kadra | 1.7 |
| T26 | Kadra | 1.6 |
| T27 | Kadra | 1.8 |
| T28 | Kadra | 1.9 |
| T29 | Kadra | 1.7 |
| T30 | Kumbarwada | 2 |
| T31 | Kumbarwada | 1.4 |
| T32 | Kumbarwada | 1.35 |
| T33 | Kumbarwada | 1.5 |
| T34 | Kumbarwada | 1.25 |
| T35 | Kumbarwada | 2 |
| T36 | Kumbarwada | 1.9 |
| T37 | Kumbarwada | 1.1 |
| T38 | Kumbarwada | 1.6 |
| T39 | Kumbarwada | 1.3 |
| T40 | Kumbarwada | 1.85 |
| T41 | Castle Rock | 1.9 |
| T42 | Castle Rock | 1.3 |
| T43 | Castle Rock | 1.7 |
| T44 | Castle Rock | 1.9 |
| T45 | Castle Rock | 2 |
| T46 | Castle Rock | 1.5 |
| T47 | Castle Rock | 2 |
| T48 | Castle Rock | 1.95 |
| T49 | Castle Rock | 1.9 |
| T50 | Castle Rock | 1.4 |
| T51 | Castle Rock | 1.35 |
| T52 | Castle Rock | 1.9 |
| T53 | Castle Rock | 1.6 |
| T54 | Phansoli | 1.6 |
| T55 | Phansoli | 1.5 |
| T56 | Phansoli | 1.4 |
| T57 | Phansoli | 2 |
| **Total** |  | **100.1** |

**Table S2.** List of tree species found in sampling plots that are food plants of hornbills, based on literature (Ahirbudhyan et al., 2026) and personal observations.

| Sl. No. | Species |
| --- | --- |
| 1 | *Aphanamixis polystachya* |
| 2 | *Aporosa cardiosperma* |
| 3 | *Artocarpus heterophyllus* |
| 4 | *Artocarpus hirsutus* |
| 5 | *Acacia auriculiformis* |
| 6 | *Beilschmiedia dalzellii* |
| 7 | *Bischofia javanica* |
| 8 | *Bridelia retusa* |
| 9 | *Bridelia stipularis* |
| 10 | *Capparis rheedei* |
| 11 | *Caryota urens* |
| 12 | *Casearia graveolens* |
| 13 | *Celtis tetrandra* |
| 14 | *Cinnamomum malabattrum* |
| 15 | *Cinnamomum sp.* |
| 16 | *Diospyros montana* |
| 17 | *Dysoxylum gotadhora* |
| 18 | *Elaeagnus conferta* |
| 19 | *Ficus drupacea* |
| 20 | *Ficus exasperata* |
| 21 | *Ficus tsjakela* |
| 22 | *Flacourtia montana* |
| 23 | *Glycosmis pentaphylla* |
| 24 | *Gnetum edule* |
| 25 | *Grewia tiliifolia* |
| 26 | *Heynea trijuga* |
| 27 | *Knema attenuata* |
| 28 | *Lannea coromandelica* |
| 29 | *Litsea coriacea* |
| 30 | *Litsea nigrescens* |
| 31 | *Macaranga peltata* |
| 32 | *Madhuca longifolia* |
| 33 | *Mimusops elengi* |
| 34 | *Myristica beddomei* |
| 35 | *Myristica malabarica* |
| 36 | *Neolitsea foliosa var. scrobiculata* |
| 37 | *Tetrapilus dioicus* |
| 38 | *Monoon fragrans* |
| 39 | *Prunus ceylanica* |
| 40 | *Sarcostigma kleinii* |
| 41 | *Schleichera oleosa* |
| 42 | *Sterculia guttata* |
| 43 | *Strychnos nux-vomica* |
| 44 | *Syzygium densiflorum* |
| 45 | *Syzygium caryophyllatum* |
| 46 | *Syzygium cumini* |
| 47 | *Syzygium hemisphericum* |
| 48 | *Syzygium rubicundum* |
| 49 | *Syzygium zeylanicum* |
| 50 | *Tabernaemontana alternifolia* |
| 51 | *Uvaria concava* |
| 52 | *Vitex altissima* |
| 53 | *Zanthoxylum rhetsa* |
| 54 | *Ziziphus oenopolia* |
| 55 | *Ziziphus rugosa* |

**Table S3.** Total number of detections (opportunistic and within transects) of Malabar Grey Hornbill, Malabar Pied Hornbill and Great Hornbill.

| Species | Detections<br>in transects | Opportunistic<br>detections | Total number<br>of detections |
| --- | --- | --- | --- |
| Malabar Grey Hornbill | 140 | 66 | 206 |
| Malabar Pied Hornbill | 13 | 102 | 115 |
| Great Hornbill | 20 | 17 | 37 |

**Table S4.** Multi-model inference table for the distance sampling models used to estimate abundance and density of Malabar Grey Hornbill in KTR. Cutpoints used—0, 30, 70, 100 and 150.

| Sl. No. | Key function | Formula | $p$-value ($\chi^2$<br>GoF) | $\hat{P}_a$ | SE ($\hat{P}_a$) | $\Delta$ AIC |
| --- | --- | --- | --- | --- | --- | --- |
| 1 | Half-normal | ~1 | 0.24 | 0.79 | 0.08 | 0 |
| 2 | Hazard-rate | ~1 | 0.13 | 0.82 | 0.11 | 1.48 |
| 3 | Half-normal | ~Range | NA | 0.78 | 2309.64 | 5.77 |
| 4 | Hazard-rate | ~Range | NA | 0.87 | 0.05 | 8.97 |

**Table S5a.** Pearson’s coefficients and *p*-values for correlation analysis of encounter rates of Malabar Grey Hornbill with habitat and resource variables. Rows in bold have *p*-values below 0.05, indicating statistically significant correlations.

| Variable | Pearson's coefficient ($r$) | $p$ |
| --- | --- | --- |
| Stem density | 0.72 | 0.07 |
| Basal area (m<sup>2</sup> ha<sup>-1</sup>) | 0.55 | 0.20 |
| Canopy height (m) | -0.51 | 0.24 |
| **Food-tree stem density (ha<sup>-1</sup>)** | **0.84** | **0.02** |

**Table S5b.** Pearson’s coefficients and *p*-values for correlation analysis of encounter rates of Malabar Pied Hornbill with habitat and resource variables. Rows in bold have *p*-values below 0.05, indicating statistically significant correlations.

| Variable | Pearson's coefficient (*r*) | *p* |
| --- | --- | --- |
| Stem density | -0.5 | 0.25 |
| **Basal area (m<sup>2</sup> ha<sup>-1</sup>)** | **-0.78** | **0.04** |
| Canopy height (m) | 0.14 | 0.77 |
| Food-tree stem density (ha<sup>-1</sup>) | -0.43 | 0.34 |

**Table S5c.** Pearson’s coefficients and *p*-values for correlation analysis of encounter rates of Great Hornbill with habitat and resource variables. Rows in bold have *p*-values below 0.05, indicating statistical significant correlations.

| Variable | Pearson's coefficient (*r*) | *p* |
| --- | --- | --- |
| Stem density | -0.4 | 0.37 |
| Basal area (m<sup>2</sup> ha<sup>-1</sup>) | -0.71 | 0.07 |
| Canopy height (m) | -0.05 | 0.92 |
| Food-tree stem density (ha<sup>-1</sup>) | -0.31 | 0.50 |

